# An Association Test of the Spatial Distribution of Rare Missense Variants within Protein Structures Improves Statistical Power of Sequencing Studies

**DOI:** 10.1101/2021.08.09.455695

**Authors:** Bowen Jin, John A. Capra, Penelope Benchek, Nicholas Wheeler, Adam C. Naj, Kara L. Hamilton-Nelson, John J. Farrell, Yuk Yee Leung, Brian Kunkle, Badri Vadarajan, Gerard D. Schellenberg, Richard Mayeux, Li-san Wang, Lindsay A. Farrer, Margaret A. Pericak-Vance, Eden R. Martin, Jonathan L. Haines, Dana C. Crawford, William S. Bush

## Abstract

Over 90% of variants are rare, and 50% of them are singletons in the Alzheimer’s Disease Sequencing Project Whole Exome Sequencing (ADSP WES) data. However, either single variant tests or unit-based tests are limited in the statistical power to detect the association between rare variants and phenotypes. To best utilize rare variants and investigate their biological effect, we exam their association with phenotypes in the context of protein. We developed a protein structure-based approach, POKEMON (Protein Optimized Kernel Evaluation of Missense Nucleotides), which evaluates rare missense variants based on their spatial distribution on the protein rather than allele frequency. The hypothesis behind this is that the three-dimensional spatial distribution of variants within a protein structure provides functional context and improves the power of association tests. POKEMON identified four candidate genes from the ADSP WES data, namely two known Alzheimer’s disease (AD) genes (*TREM2* and *SORL*) and two novel genes (*DUSP18* and *CSF1R*). For known AD genes, the signal from the spatial cluster is stable even if we exclude known AD risk variants, indicating the presence of additional low frequency risk variants within these genes. *DUSP18* has a cluster of variants primarily shared by case subjects around the ligand-binding domain, and this cluster is further validated in a replication dataset with a larger sample size. POKEMON is an open-source tool available at https://github.com/bushlab-genomics/POKEMON.

## INTRODUCTION

High-throughput DNA sequencing of diverse human populations has identified millions of genetic variants, the vast majority of which are exceptionally rare. A survey of ~60,000 individuals from the Exome Aggregation Consortium (ExAC) found that out of ~7M variants, 99% have a frequency <1% and 54% are singletons (Karczewski et al., 2020). Similarly, in the Alzheimer’s Disease Sequencing Project (ADSP) Whole Exome Sequencing (WES) of ~10k individuals, 97% of identified variants have a minor allele frequency <1% and 23% are singletons (Butkiewicz et al., 2018). However, the effect of most rare variants on diseases of interest remains unknown because of insufficient statistical power to detect the associations between these variants and phenotypes.

We hypothesized that rare variants contribute to common diseases by forming clustered or dispersed patterns within protein structures that reflect modest disruption of protein function. Based on this hypothesis, incorporating protein spatial context should improve rare variant association tests. Prior studies have shown missense variants exhibit non-random patterns in protein structures, such as cancer-associated hot spot regions with a high density of missense somatic mutation (Tokheim et al., 2016). Our group (Sivley et al., 2018) also found that germline causal missense variants for Mendelian diseases exhibit non-random patterns in 3D space, including both clusters and depletion.

To test this hypothesis, we developed a kernel function to quantify genetic similarity among individuals by using protein structure information. Consider a scenario where two individuals have different missense variants distal in genomic coordinates but close in 3D protein structure; these individuals will be assigned a high genetic similarity through our kernel function, which when applied over an entire dataset captures the spatial patterns of rare missense variants. Using a statistical framework similar to SKAT (Wu et al., 2011), we test the association of rare variants with quantitative and dichotomous phenotypes using this structure-based kernel. We call this approach POKEMON (Protein Optimized Kernel Evaluation of Missense Nucleotides). We validated that POKEMON can identify trait associations with spatial patterns formed by missense variants both in simulation studies and real-world data.

POKEMON identified four candidate genes from the ADSP WES Discovery Dataset, namely two AD genes (*TREM2* and *SORL1*) and two novel genes (*DUSP18* and *CSF1R*). Both *DUSP18* and *CSF1R* have clusters of variants primarily shared by case subjects around ligand-binding sites. Specifically for the cluster identified in *DUSP18*, we examined it in the ADSP WES replication dataset with a larger sample size and found that the cluster is better formed by the additional variants included. In summary, the cluster we identified is populated with variants mostly from cases and likely has a functional association with AD risk.

By performing an association test in the context of protein structure, we have identified highly relevant gene-disease associations which are driven by specific clusters of variants. Such clusters imply functional domains in the protein structure susceptible to variation and related to disease risk. Analyzing missense variants from complex disease studies in this way provides a new structural aspect which can be leveraged for association tests.

## RESULTS

### POKEMON can detect associations with spatially clustered or dispersed rare variants

As a proof of concept, we evaluated the performance of POKEMON using simulations that mimic real-world case/control studies. The simulation datasets varied in sample sizes (1000 - 5000), the odds ratio of the core variants (2.0 - 3.0), and the proportion of influential to neutral variants (0.3 - 0.9). For a cluster pattern, influential variants were simulated by establishing a maximum odds ratio decaying over a fixed distance of14 Å. We limit the number of variants within the genotype profile to 50, which is the mean number of variants mapped per protein in the ADSP WES. For each simulation scenario, we generated 100 datasets and tested for spatial clustering using a structure kernel, estimating power as the proportion of significant tests. In general, POKEMON performed better than two other structure-based methods, PSCAN(Tang et al., 2020) and POINT(Marceau West et al., 2019) and a frequency-based kernel (SKAT) (see Figure 1). For scenarios with weaker effects and/or smaller sample sizes, POKEMON showed much better performance relative to other methods.

**Figure 1.**
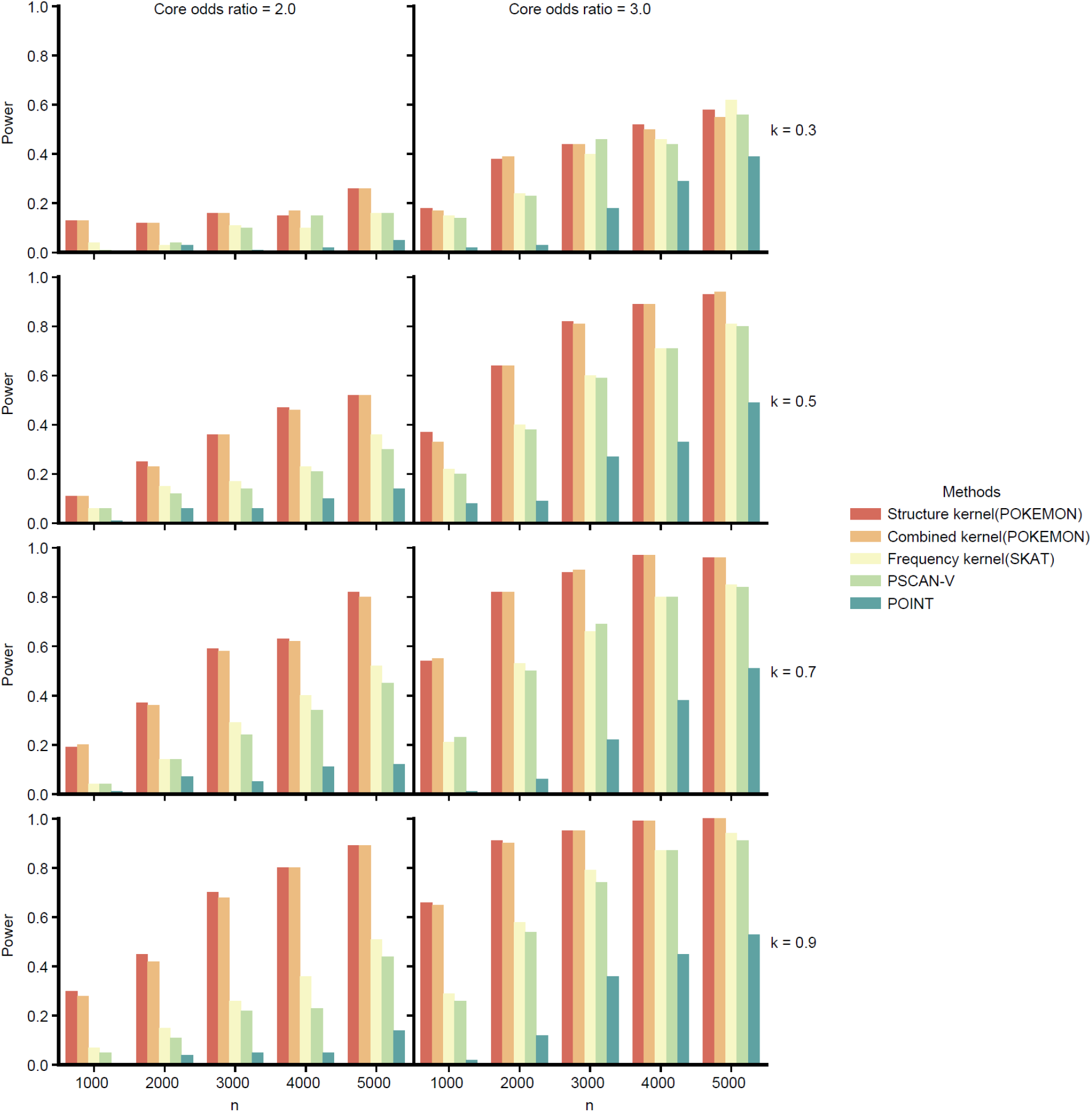
Empirical power to detect a pattern of association centered at the protein core among five methods with varying percentages of influential variants (k) and core variant odds ratios. Empirical power is calculated by counting the number of tests with a p-value below the significance level in 100 independent tests. The five methods include the structure kernel (POKEMON), combined kernel (POKEMON), frequency kernel (SKAT), PSCAN with variance test (PSCAN-V), and POINT. The core variant odds ratio is chosen as 2.0 or 3.0 (left to right). *k* is the percentage of pathological variants within the selected 50 variants, ranging from 0.3 to 0.9 (from top to bottom).

Additionally, to evaluate POKEMON’s ability to identify a dispersed pattern, we simulated the scenario where variants are distributed on the protein’s surface. When all influential variants on the surface had small odds ratios, none of the methods performed well. When increasing the odds ratio to 1.5, POKEMON outperformed other methods in most scenarios, except for cases where the percentage of influential variants was low (0.3) (see Figure S1).

We also assessed POKEMON’s power at a higher resolution for different configurations of core odds ratios and influential variant proportions. Figure 2 illustrates the dynamics of statistical power for the POKEMON test under the assumption of a spatial effect. POKEMON achieved a power of 0.8 with study designs commonly found in sequencing studies of complex disease: a population of 3000 cases/3000 controls, the core odds ratio of 3.0, and 50% of the rare variants influential on the simulated phenotype with moderate effect. However, as expected, when the percentage of influential variants is low (<35%) and the core variant odds ratio is small (<1.8), POKEMON did not reach 80% power. A small core odds ratio and a low percentage of influential variants are more challenging for POKEMON to assess because more control subjects will carry variants located within the cluster region, making POKEMON less likely to identify associated patterns.

**Figure 2.**
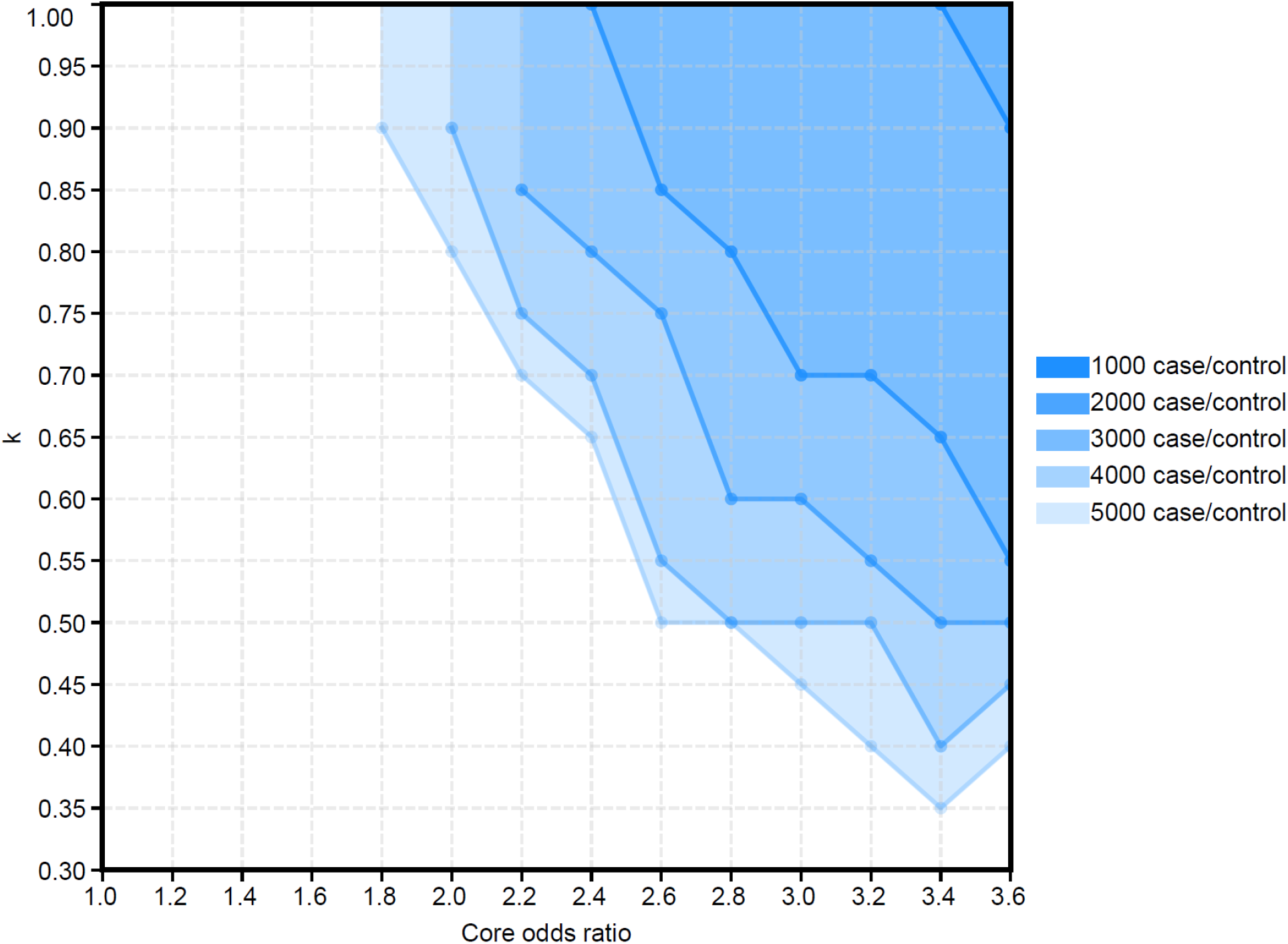
Power assessment for POKEMON at different configurations (for both structure kernel and combined kernel). Each line represents the minimum percentages of influential variants (k) and minimum core variant odds ratios to reach a power of 0.8 when the number of cases/controls is fixed.

### POKEMON replicates the cancer-related spatial clusters from the TCGA dataset

To demonstrate POKEMON’s ability to identify spatial patterns from real-world data, we analyzed germline variants from The Cancer Genome Atlas (TCGA), which has previously been evaluated for spatial clusters associated with cancer risk and metastasis (Mashl et al., 2018). We constructed a case/control dataset by combining 10389 subjects from TCGA across 33 cancer types with 4919 presumably cancer-free controls from the ADSP WES Discovery Dataset. We restricted our POKEMON analysis to rare variants with unknown effects and 31 proteins with functional assessement in the literature. This analysis directly tested the hypothesis that cancer-related variants tend to cluster in a protein hotspot while rare variants from cancer-free subjects are randomly distributed. We observed significant enrichment of statistical associations within the 31 proteins evaluated (27 with p < 0.05, see Supplementary Table 1).

From these results, we focus specifically on two genes highlighted in the literature that have been experimentally validated, namely *RET* and *MET* (Table 1). We found similar patterns of variant clustering for *RET* and *MET*, formed by somatic variants and pathological/likely pathological germline variants (Mashl et al., 2018). For *MET*, POKEMON identified a cluster formed by P1091L, S1092G, and Q1085K, which surrounds the pathological variant H1112R (Figure 3A and B). For *RET*, POKEMON identified a cluster formed by E867D, V871I, L895F, R897L, and R908K, which surrounds the pathological variants R912P and R918T (Figure 3C and D). Notably, POKEMON identified these two clusters via case/control analysis of rare germline variants while excluding known pathological variants. Thus our significant association statistic is driven by additional rare variants within *MET* and *RET* surrounding those with known pathological effects.

**Figure 3.**
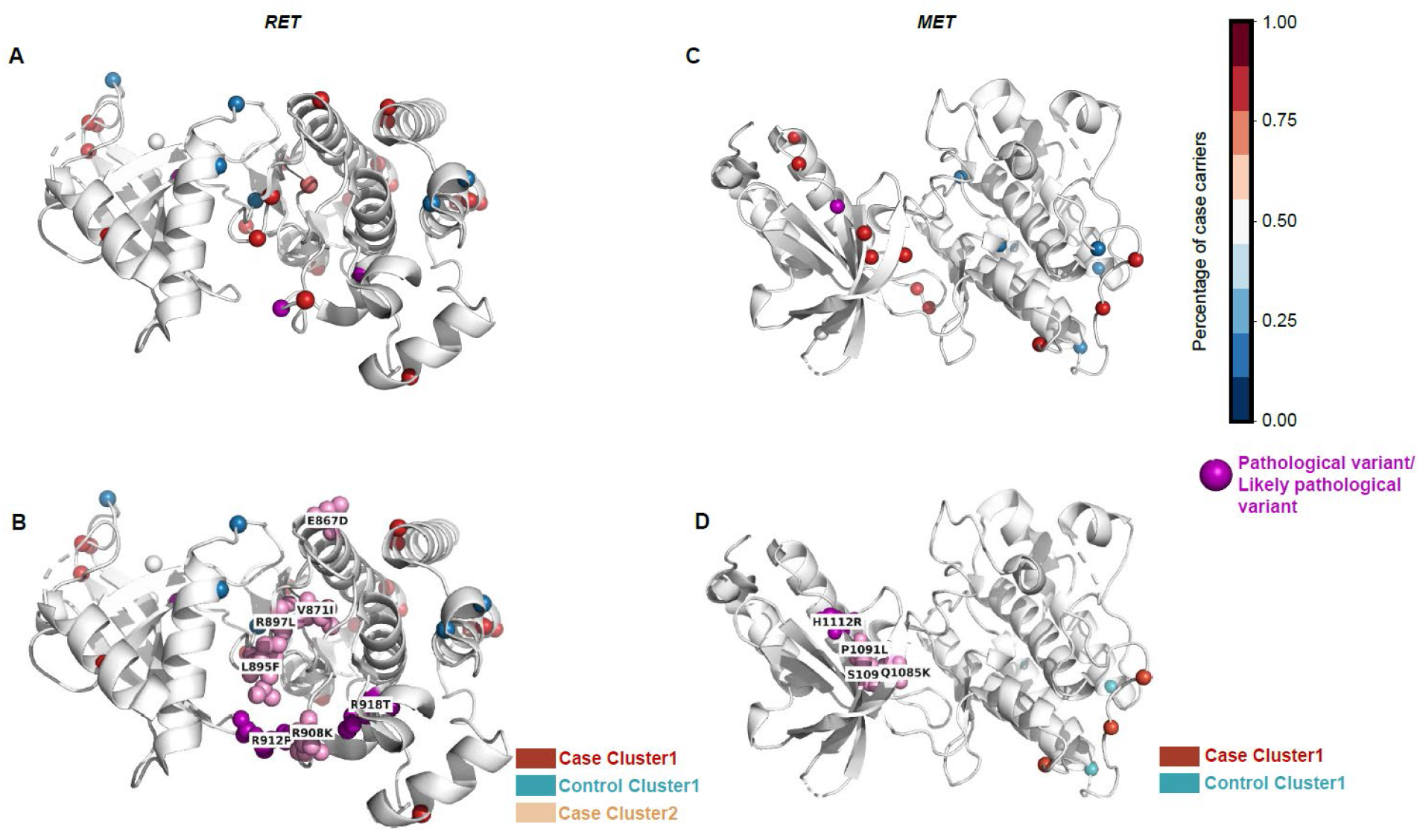
Spatial distribution of variants from TCGA dataset within MET (PDB:1R0P) and RET (PDB:2IVT). A and C show the rare missense variants with unknown effects mapped to the structure. The color scale indicates the percentage of case subjects that carry the variants out of the overall sample. Pathological and likely pathological variants are highlighted in purple. B, D show clusters identified by POKEMON. Clusters that are consistent with the original literature are highlighted with pink sphere models. Pathological and likely pathological variants are highlighted with purple sphere models.

**Table 1:** Results for *MET and RET* from TCGA dataset

| Gene | PDB entry | Phenotype | # SNPs | #SNPs mapped | p-value |
| --- | --- | --- | --- | --- | --- |
| *MET* | PDB:1R0P | cancer | 757 | 17 | 5.454e-04 |
| *RET* | PDB:2IVT | cancer | 765 | 33 | 1.606e-04 |

**Table 2:** genes associated with AD based on structure kernel

| Gene | # SNP | # SNP mapped | PDB Entry | p-value | log(p-value) |
| --- | --- | --- | --- | --- | --- |
| *TREM2* | 40 | 33 | PDB:6XDS | 1.261E-08 | 7.899 |
| *SORL1* | 203 | 56 | PDB:3WSY | 8.167E-05 | 4.088 |
| *CSF1R* | 74 | 38 | PDB:4LIQ | 4.428E-04 | 3.354 |
| *DUSP18* | 25 | 18 | PDB:2ESB | 6.717E-04 | 3.509 |

### POKEMON identifies known AD risk genes (*TREM2* and *SORL1*) and novel candidate genes (*CSF1R and DUSP18*)

To discover any spatial rare variant patterns associated with AD, we applied POKEMON with a structure kernel to the ADSP WES Discovery Dataset with 5,522 AD cases and 4,919 controls. We perform the POKEMON test on 4,173 genes with available protein structures and ≥ 5 rare missense variants (MAF<0.05). *APOE* ε2 and ε4 dosages were included as covariates.

We used two significance thresholds to identify candidate genes: a typical Bonferroni correction threshold and an empirical threshold we derived from the *MET* and *RET* results based on our TCGA analysis which reflects the order of magnitude of significance of a true-positive signal obtained from variants with unknown effects and extremely low allele frequency (<0.001).

Overall, there are four genes out of 4,173 identified as candidate genes. *TREM2* was identified with the Bonferroni correction, while *SORL1, CSF1R*, and *DUSP18* were identified with the empirical threshold (<1E-03).

#### The spatial cluster is stable for *SORL1*, while *TREM2* is driven primarily by a single variant

As with our TCGA analyses of *RET* and *MET*, both *SORL1* and *TREM2* harbor known AD-assocated variants. To determine if the cluster pattern we detected is stable even in the absence of these known effects, we excluded any AD-related variants previously identified in GWAS studies leaving only rare genetic variants with unknown effects within *SORL1* and *TREM2*. A significant result from this analysis indicates that rare additional variants within these genes contribute to AD risk.

Indeed, for *SORL1*(Ensembl: ENST00000260197; PDB: 3WSY), even though AD-related variants A528T (Overall MAC:439; MAF:0.0210), E270K (Overall MAC:990; MAF:0.0474), and T947M (Overall MAC:2; MAF: 9.578e-05) were excluded respectively (Vardarajan et al., 2015), the signals persist. While the results for the structure kernel with these loci excluded are still significant, *SORL1* is not significantly associated with AD even with all variants included for the frequency kernel analysis (Table 3a). The result indicates that the spatial pattern of variants within the 3WSY structure of *SORL1* is associated with AD.

**Table 3a:** Results for *SORL1* w/ and w/o known loci

| Method | Gene | PDB Entry | p-value |
| --- | --- | --- | --- |
| Structure kernel | SORL1 | PDB:3WSY | 8.167E-05 |
|  | SORL1(exclude A528T) |  | 2.841E-02 |
| Frequency kernel (SKAT) | SORL1 |  | 0.0923 |
|  | SORL1(exclude A528T) |  | 8.578E-01 |
| Structure kernel | SORL1 | PDB:3WSY | 8.167E-05 |
|  | SORL1(exclude E270K) |  | 7.373E-05 |
| Frequency kernel (SKAT) | SORL1 |  | 0.0923 |
|  | SORL1(exclude E270K) |  | 3.881E-02 |
| Structure kernel | SORL1 | PDB:3WSY | 8.167E-05 |
|  | SORL1(exclude T947M) |  | 8.166E-05 |
| Frequency kernel (SKAT) | SORL1 |  | 0.0923 |
|  | SORL1(exclude T947M) |  | 9.214E-02 |

**Table 3b:**
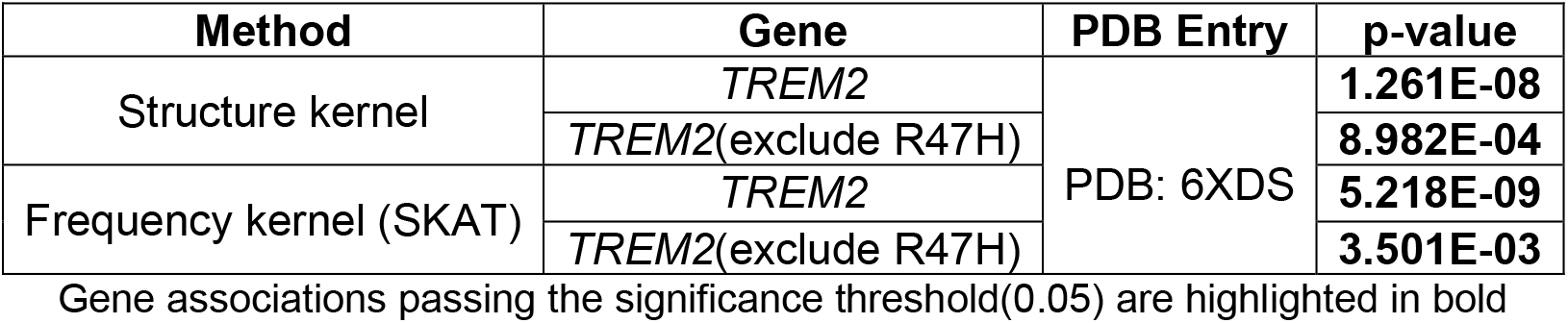
Results for *TREM2* w/ and w/o known loci

For *TREM2* (Ensembl: ENST00000373113; PDB: 5ELI), The signal is likely driven primarily by the variant R47H. The results for *TREM2* from the structure kernel test are comparable to that from the frequency kernel test before and after R47H being excluded. Excluding R47H (Korvatska et al., 2015) changes the p-value drastically for both the structure kernel and the frequency kernel (Table 3b).

#### *DUSP18* has a cluster of variants primarily shared by case subjects around the ligand-binding domain

The signal identified within *DUSP18* (Ensembl: ENST00000334679; PDB: 2ESB) is driven by a cluster of variants primarily shared by case subjects in the catalytic domain for ligand binding, highlighted as Case Cluster 1 in Figure 4. This association is strengthened in the replication dataset (Table 4), where the cluster is better formed by an additional case variant A35T and the deletion of a control variant G107A (Figure 4). Overall, Case Cluster 1 is formed by missense variants A35T, N51S, E55G, S74A, R78C, and R110H, which surround the ligand-binding site (Figure 5). Indeed, Variant R110H, a catalytic triad residue, interacts with E55G (Jeong et al., 2006), indicating a functional interaction related to AD risk. *DUSP18* has been previously reported to inhibit SUMOylation and reduce *ATXN1* aggregation (Ryu and Lee, 2018), and *ATXN1* loss of function is associated with an increased risk for AD (Suh et al., 2019).

**Figure 4.**
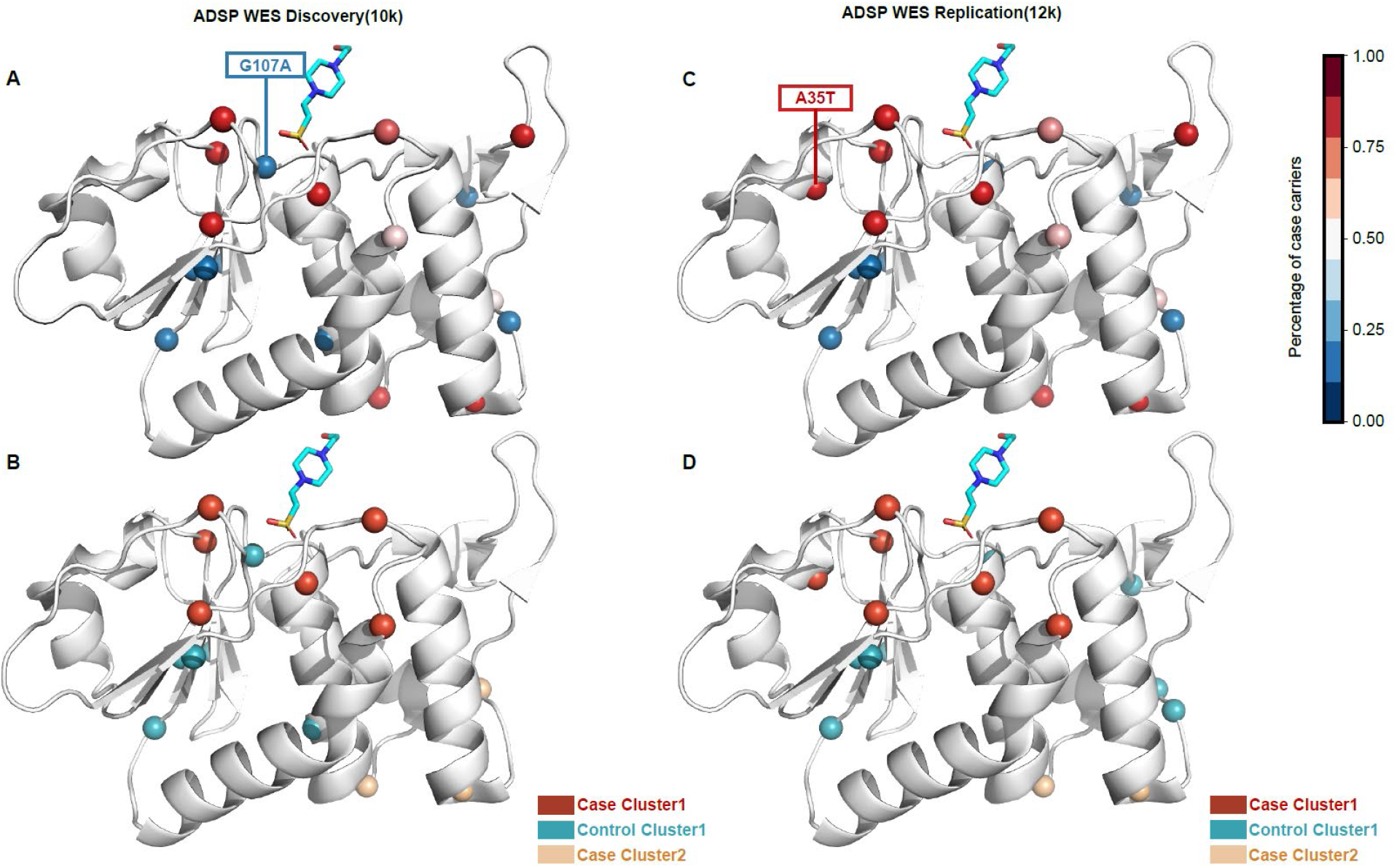
Spatial distribution of variants from ADSP WES discovery and replication dataset within DUSP18 (PDB:2ESB.A). A and C show the rare missense variants with unknown effects mapped to the structure. The color scale indicates the percentage of case subjects that carry the variants out of the overall sample. Variants that show a different percentage of case carriers and are being classified into other clusters between ADSP WES discovery and replication dataset are highlighted and labeled. B, D show clusters identified by POKEMON. Clusters are colored differently.

**Figure 5.**
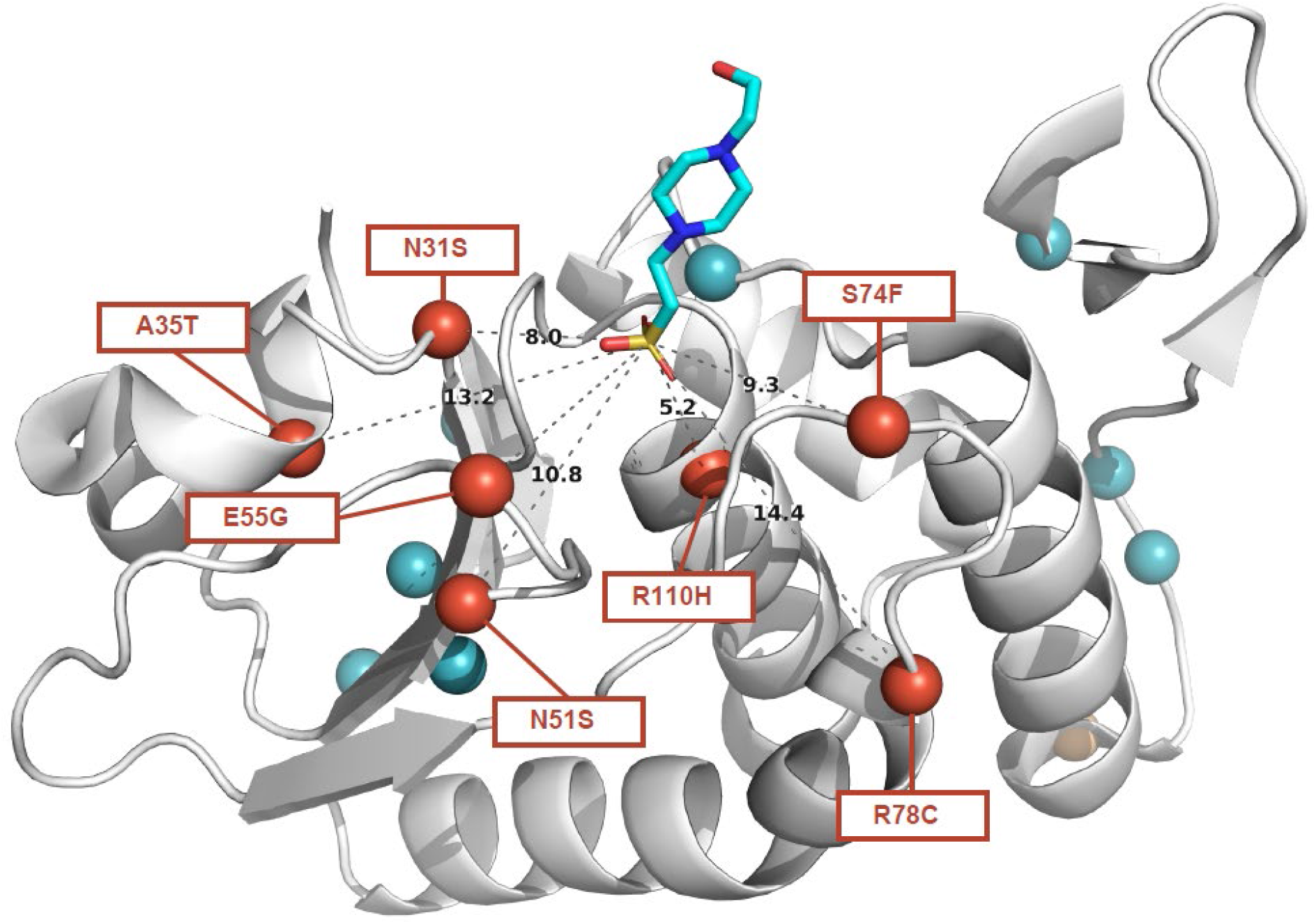
Spatial distribution of variants and their distance to the EPE within DUSP18 (PDB:2ESB.A). The distance is calculated from Cα to S. All the variants that belong to the cluster of interest are highlighted and labeled with the AA change.

**Table 4:** Results for candidate genes from the replication dataset

|  |  | **ADSP WES Discovery(10k)**<br>Overall: 10441<br>Case/Control: 5522/4919 |  |  | **ADSP WES replication (12k)**<br>Overall: 11828<br>Case/Control: 6238/5590 |  |  |
| --- | --- | --- | --- | --- | --- | --- | --- |
| **Gene** | **PDB entry** | **# SNPs** | **#SNPs mapped** | **p-value** | **# SNPs** | **#SNPs mapped** | **p-value** |
| *TREM2* | PDB:6XDS | 40 | 33 | 1.261E-08 | 61 | 29 | 5.049E-11 |
| *SORL1* | PDB:3WSY | 203 | 56 | 8.167E-05 | 446 | 59 | 9.754E-06 |
| *CSF1R* | PDB:4LIQ | 74 | 38 | 4.428E-04 | 196 | 38 | 6.128E-03 |
| *DUSP18* | PDB:2ESB | 25 | 18 | 6.717E-04 | 37 | 19 | 1.037E-05 |

#### The spatial pattern within *CSF1R* is unstable through the replication dataset

For *CSF1R* (Ensembl: ENST00000286301; PDB: 4LIQ), the structure kernel result is more significant than the frequency kernel (Table 5), indicating that spatial patterns within the protein improve detection power. A case cluster identified by POKEMON in the Discovery Dataset contains T163A, N153H, A123G, L111I, R106Q, R106W, Q77E, and P53L (Figure 6.B: Case Cluster 1). The distances between those variants range from 5.3 to 14.3 Å, while their average sequence distance is 18 amino acids. This cluster is within the extracellular region (AA 1-498), where CSF-1 or IL-34 binds (Stanley and Chitu, 2014).

**Table 5:**
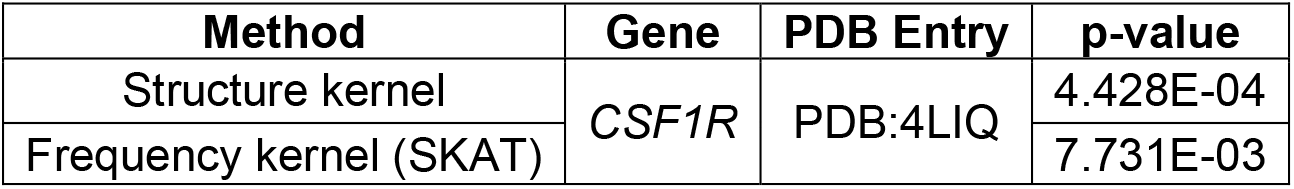
Results for *CSF1R* with different kernels

| Method | Gene | PDB Entry | p-value |
| --- | --- | --- | --- |
| Structure kernel | *CSF1R* | PDB:4LIQ | 4.428E-04 |
| Frequency kernel (SKAT) |  |  | 7.731E-03 |

**Figure 6.**
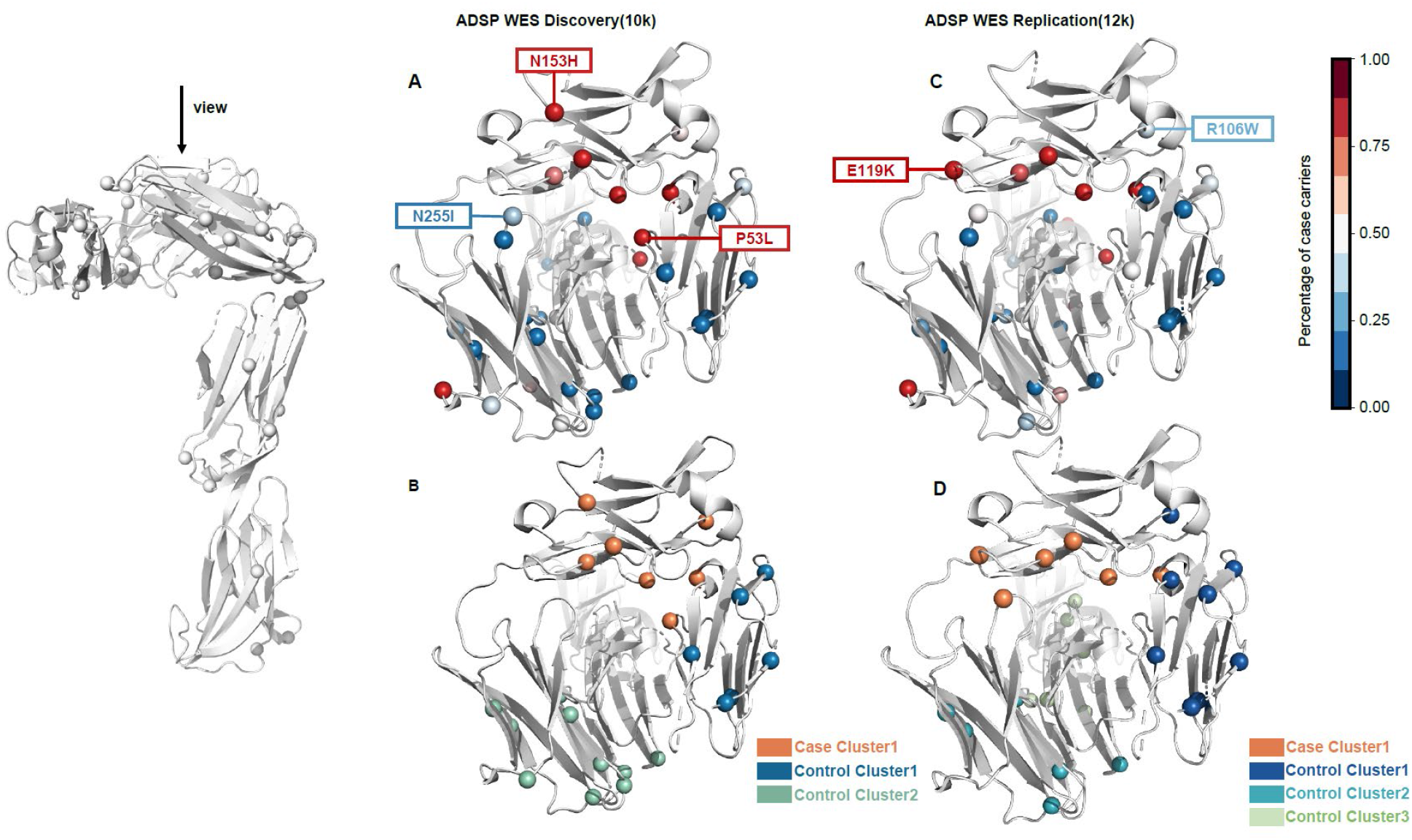
Spatial distribution of variants from ADSP WES discovery and replication dataset within CSF1R (PDB:4LIQ.A+E). A and C show the rare missense variants with unknown effects mapped to the structure. The color scale indicates the percentage of case subjects that carry variants from the overall sample. Variants that show a different percentage of case carriers and are classified into other clusters (illustrated by color) between ADSP WES discovery and replication dataset are highlighted and labeled. B, D show clusters identified by POKEMON.

However, when we examined *CSF1R* with the replication dataset, we did not observe an improved signal compared to the Discovery Dataset (Table 4). Reviewing the PDB:4LIQ structure, we found that Case Cluster 1 persists in the replication dataset while other clusters become sporadic. Specifically, Case Cluster 1 has two additional case variants E119K and N255I, while two case variants P53L, N153H drop out of the dataset (Figure 6.A, C). The complete comparison of variants between the Discovery Dataset and the Replication dataset can be found in the Supplementary Tables.

## Discussion

We have shown that POKEMON improves the power to detect rare variant associations in the context of protein structure. We found POKEMON outperforms other structure-based methods through simulation studies except in a small number of cases where all existing methods have insufficient power. Specifically, POKEMON achieves a power of 0.8 for what we presume to be a common scenario in sequencing studies: a study population of 3,000 cases/3,000 controls, a core odds ratio of 3, and 50% of the variants associated with the phenotype. In contrast, all other methods tested for this scenario have power below 0.6. We applied POKEMON to the ADSP Discovery WES dataset and identified spatial patterns of rare variants related to AD risk. The spatial patterns of variants within *SORL1*, is consistent with a previously reported association using PSCAN(Tang et al., 2020). We also identify two potentially AD-associated clusters of variants within *CSF1R* and *DUSP18*, located around ligand-binding sites. Specifically, the cluster within *DUSP18* is validated with the replication dataset with a larger sample size.

Notably, an advantage for POKEMON over other rare-variant analysis methods is that statistical power increases with the observation of any new variant including singletons assuming the existence of divergent spatial patterns between cases and controls. In most rare variant association tests, increasing sample size only increases the power for non-singleton variants in the resulting data. Even for those non-singleton variants, the improvement in power is not necessarily proportional to the increase in sample size. Moreover, additional neutral variants will be introduced and negatively impacting the statistical power when the sample size increases. In contrast, POKEMON can utilize rare variants and even singletons with the structure kernel, regardless of their low allele frequency. The increasing number of rare variants helps form a spatial pattern, which can be identified by POKEMON with higher power (Figure S4D).

Based on our analyses of *MET/RET* in the TCGA dataset, and SORL1 in the ADSP WES Discovery dataset, we also demonstrated that the association detected by our spatial kernel is not driven by a single variant but rather a collection of variants with modest effects. Additionally, even though we didn’t include any population-related covariates in the structure kernel test for ADSP WES Discovery Dataset, the overall results didn’t show large genomic inflation (GC=1.23). This confirms our assumption that the structure kernel in POKEMON is less susceptible to population stratification than frequency-based tests; any constraint to the positions of rare variants within protein structures is likely independent of the variants’ population origin and therefore does not confound analyses as is typical of a frequency-based test.

POKEMON is designed to leverage pre-existing biological information for sequencing datasets where typically only variant counts or frequencies are considered. Even though protein structure information of variants has been incorporated into association tests like POINT and PSCAN (Marceau West et al., 2019; Tang et al., 2020), they serve as guiding information for more traditional association tests ultimately based on allele frequency. Therefore, these approaches are still potentially subject to the limitations in unit-based or single variant tests. With the structure kernel, POKEMON uses the spatial information of a missense variant, which is independent of allele frequency. Assuming the rare variants form spatial patterns, POKEMON ameliorates the power dilemma induced by increasing numbers of singleton variants as the sample size of sequencing studies increases.

Importantly, POKEMON does not have sufficient power to detect patterns for some scenarios. As shown in the simulations, POKEMON lacks the power even with 5,000/5,000 cases/controls when the percentage of influential variants is low (<30%) and the core variant odds ratio is small (<1.5). While we expected lower power in this scenario, this result may also reflect a limitation of our simulation strategy; we limit the number of variants simulated to 50. If more variants are included in the test, the underlying spatial pattern is better formed, and thus the power of POKEMON will improve.

We anticipate POKEMON will be helpful as a large-scale screening method to detect potentially disease-associated proteins in a proteome-wide fashion. Currently, the PDB structures deposited in the PDBank only cover ~70% of the identified molecular functions in the human genome (Somody et al., 2017). We expect that the improvement in Cryo-EM and prediction methods like AlphaFold2 (Senior et al., 2020) will massively increase the availability of structural information for proteins and complexes. As POKEMON is a unit-based test, it only provides a single association statistic for the influence of all missense variants within the protein for a phenotype. Follow-up analyses to assess specific SNVs or refine SNV subsets may provide more detailed quantitative assessments of specific variant spatial patterns.

## MATERIAL AND METHODS

### Derivation of the POKEMON method

We briefly review the linear mixed model used in association tests and then introduce the construction of a structure kernel for POKEMON. Assume we have *n* individuals for whom we have *p* non-genetic covariates, genotypes for *m* SNPs, and the phenotype *y* as an *n* × 1 vector. Genotype ***G*** is a *n* × *m* matrix. Covariate ***X*** is a *n* × *p* matrix.

A linear mixed model contains a fixed effect from covariates ***X**β*, a random effect annotated by ***Z**u*, and an error term *ϵ*. The *y* is fit with a high-dimensional normal distribution (2). The random effect can be further divided into two parts, an environmental effect 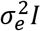 and a genetic effect 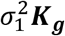. ***K**_g_* is the kernel containing the genetic similarity between individuals. 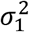 is the amount of variance of *y* explained by ***K**_g_*.

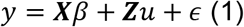

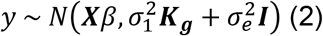

The null hypothesis *σ*_1_ = 0 indicates that ***K**_g_* does not explain any variance of *y*. The score statistic *Q* is defined as the partial differential for the log-likelihood on 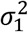. Under the null hypothesis, *Q* follows a mixed chi-squared distribution(3), where ***S*** projects *y* into a space orthogonal to covariates and *λ_i_* are the eigenvalues of ***SK**_g_**S***.

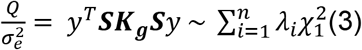

For POKEMON, we construct the *n* × *n* kernel ***K**_g_* in the context of protein as follows. For ***K**_g_*, each entry is the genetic similarity between individuals based on the variants they carry, which is weighted by variants’ distance in the protein structure(4). *d_kl_* is the distance of pair-wise single nucleotide variants (SNVs) in angstroms (Å) within the protein. *k* and *l* represent the *k*th variant from individual *i* and the *lth* variants from individual *j*.

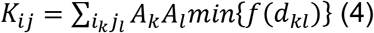

Some protein structures are formed by identical subunits (homo-multimer), which introduces redundancy in the variant-to-amino acid projection (i.e., one variant can map to multiple amino acids located in different subunits). To eliminate the spatial similarity induced by multiple mapping locations of a single variant in a homo-multimer, we took *d_kl_* to be the minimum distance over all pair-wise distances. Function *f(d)* converts a Euclidean distance to the similarity score for a pair of variants.

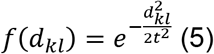

As a default, the exponential function for *f* in (5) with *t* set to a value of 7 Å, with which the effect decays below 0.1 after 2t (14 Å). 14 Å was chosen as it is a commonly adopted short-range non-bonded cut-off in molecular dynamic simulation (Monticelli et al., 2008).

Apart from spatial patterns, we also account for the magnitude of the protein change due to the different amino acid substitutions. We scaled the pair-wise variants by their amino acid substitution, which is defined as *A_k_* and *A_l_*. *A_k_* and *A_l_* are the weights for amino acid substitution for variant *k* and variant *l* according to the BLOSUM62 matrix (Henikoff and Henikoff, 1992), respectively. For a less conservative amino acid substitution, the score *s_k_* in BLOSUM62 matrix will be negative and consequently *A_k_* will be greater than 1. In contrast, for a neutral or conservative amino acid substitution, *s_k_* will be positive and *A_k_* will be less than 1.

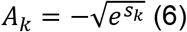

The structure kernel is nonlinear in contrast to the SKAT tests (Wu et al., 2011), which uses a linear kernel (e.g., ***K*** = ***GWW’G’***) to calculate the genetic similarity between individuals. The genetic similarity in a linear kernel between individuals is the sum of weighted SNVs being shared. However, singletons are carried by only a single individual and thus fail to be included in calculating genetic similarity. With the structure kernel, a pair of singleton variants will be assigned non-zero weights if they are spatially proximate in the protein structure. The interpretation of the structure kernel is that case individuals are genetically similar because they share more spatially clustered or dispersed rare variants than the control individuals.

We also allow for incorporating allele frequency in the POKEMON test and develop a combined kernel function. Variants clustered in protein structure already contribute to a high genetic similarity based on our structure kernel. With a *combined* kernel, those variants will be further up-weighted if they are rare in allele frequency and vice versa. The combined kernel function is based on ***K**_g_*, extended by further scaling variants by weights derived from the allele frequency. *w_k_* = Beta(*MAF_k_; a, b*) is the weight for the *k^th^* variant characterized by beta density with *a* = 1 and *b* = 25 as default.

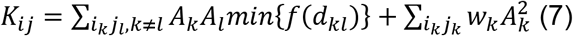

### Workflow of POKEMON

POKEMON requires a genotype matrix and consequence profile containing variant-to-amino acid mapping information as inputs (see Figure S2). Additional covariate files are optional to adjust for covariates. POKEMON first maps the variants using their coordinates into the 3D protein, which is accomplished with the consequence profile generated by Ensembl Variant Effect Predictor (VEP v95) and the reference from SIFTS function mapping a PDB entry to a UniProt residue level (Dana et al., 2019). A single variant may be mapped to multiple amino acids for multimers with identical subunits. The protein structures are fetched from the PDB during the analysis. If multiple protein structures are available for a single gene, the structure with the most variants mapped will be selected. The user may also specify a specific PDB entry. After mapping, the score between a pair of variants is calculated based on the minimum distance between them, which is further scaled by the amino acid substitution weight from the BLOSUM62 matrix by default. The pair-wise genetic similarity between individuals is the summation of all pair-wise scores of variants. The genetic similarity kernel ***K**_g_* is then evaluated in the variance component test.

### Data Simulation

We conducted simulation studies to assess POKEMON’s power in detecting disease-associated protein variant patterns. We hypothesized that variants with moderate effects on a phenotype form spatial patterns within a protein structure and induce subtle alterations to the protein’s function. To test the hypothesis, we established two patterns. The first pattern entails an embedded core within the protein disrupted by rare variants (i.e., variant clustering), while the other represents the localization of influential variants to the protein’s surface (i.e., variant dispersion). Both patterns are shown in Fig S4.A & Fig S4.B. We randomly selected a protein structure, Human c-Fms Kinase Domain (PDB:2OGV), to carry out simulations. Structural information for PDB:2OGV is available for both PSCAN and SKAT.

We simulated a clustering pattern by distributing influential variants within the core of the protein structure and scaling the variant odds ratios proportionally to their distance from the core. We then randomly sampled 50 variants from the protein. The minor allele frequencies for all the variants were randomly sampled from a log-transformed uniform distribution within an interval (−4, −2.3). This variant sampling strategy restricted the selected minor allele frequencies within the range (0.0001, 0.005) and generates singletons, which is consistent with ADSP WES studies (Figure S3). To investigate how neutral variants influence the power, we varied the percentage of influential variants out of all variants being sampled (Figure S4). For each set of parameters (e.g., sample size, core variant odds ratio, etc.), the empirical power was estimated by the percentage of successful tests out of 100 independent tests with a significance level of 0.05. We compared the empirical power of POKEMON with three other methods: SKAT, PSCAN-V, and POINT. The number of case and control subjects sampled is from 1000 to 5000. Additional details for the simulation can be found in Figure S4 and supplementary materials.

We also simulated a dispersion pattern by distributing influential variants on the protein’s surface. Considering the selected protein PDB:2OGV is about 40 Å in diameter, we defined the surface variants as those more than 21 Å away from the core, which yielded 33 variants. All the surface variants were assigned with the same odds ratio (e.g., 1.1), while the rest were considered neutral with an odds ratio of 1. The simulation settings were similar to the clustering pattern, with the only difference that we sampled 30 variants from the protein, which allowed us to tune the percentage of influential variants to as large as 90%.

### Applying POKEMON to ADSP WES data

#### Discovery Dataset

We used the whole-exome sequencing (WES) data from the Discovery Case-Control study under the Alzheimer’s Disease Sequencing Project (ADSP). ADSP WES data contains 5,740 late-onset AD cases and 5,096 cognitively normal controls primarily of European ancestry, with 218 cases and 177 controls of Caribbean Hispanic ancestry. Cases were determined based on diagnosis using cognitive testing data and medical records, while controls were determined on their low risk of developing AD by age 85 years (Bis et al., 2018; Beecham et al., 2017). ADSP WES data is available by applying to the NIAGADS Data Sharing Service.

We selected 10,441 subjects of European ancestry from the ADSP as the study group (5,522 late-onset AD cases and 4,919 cognitively normal controls). The whole-exome sequencing for these 10,441 subjects provided ~850,000 variants, of which 97.5% have a minor allele frequency < 0.01. We retained the missense variants with minor allele frequency < 0.05 for our assessment. Overall, we selected 4,173 genes with experimentally determined protein structures and ≥ 5 rare missense variants mapped to the structure. The mean number of rare missense variants mapped per gene was ~50. To exclude signals induced by the well-known *APOE* association, we included *APOE* ε2 and ε4 dosages as covariates.

#### Replication Dataset

The ADSP FUS contains the 9389 subjects from the Discovery Phase Case-Control study plus additional 1044 cases and 1395 controls for a total of 11828 non-Hispanic white subjects. The WES data in FUS was reprocessed using joint genotype calling approaches implemented in the VCPA pipeline (Leung et al., 2019). The genotype calling approaches for the replication dataset were updated from the ATLAS genotype calling process implemented for the Discovery Dataset. Therefore, we consider that this replication dataset is a validation by expanding the sample size and accounting for variability in the variant calling process. If the spatial pattern POKEMON identified is stable, the additional variants identified in the replication dataset should also contribute to the association signal.

Our primary purpose for using the replication dataset was to validate the spatial patterns identified from the discovery analysis, there for we examined only *SORL1, TREM2, DUSP18*, and *CSF1R* genes. Tests on the replication dataset were conducted similarly to the discovery analyses, with *APOE* ε2 and ε4 dosages included as covariates to regress out the *APOE* association.

### Applying POKEMON to TCGA data

The TCGA data is a real-world, true-positive example of spatial patterns of missense variants associated with phenotypes (Kamburov et al., 2015). To create a dataset in the form of a case-control study, we combined 4919 control subjects from the ADSP WES Discovery Dataset and 10389 subjects from TCGA data diagnosed with 33 cancer types (Mashl et al., 2018). We assumed that 4919 control subjects from the ADSP WES Discovery Dataset are cancer-free controls. While this is not an ideal study design, any violation of this assumption would reduce statistical power rather than identifying spurious associations. The combined case/control dataset provided a real-world assessment of our hypothesis that rare variants from cancer tissues would form spatial patterns, while the rare variants from control subjects would be randomly distributed within the protein.

All previously identified pathological or likely pathological variants were excluded from the TCGA data. Moreover, we set a stringent MAF threshold as <0.001 to retain rare variants. In summary, the entire test was carried out to examine rare variants with unknown effects. We carried out POKEMON tests on 31 genes with potential hotspots (Mashl et al., 2018) and available protein structures. We only applied structure kernel tests with no covariate included.

## CODE AVAILABILITY

The code for this study is available at GitHub: https://github.com/bushlab-genomics/POKEMON with open access.

## DATA ACCESS

Whole Exome Sequencing Data from the Alzheimer’s Disease Sequencing Project are available via NIAGADS (NG00067 - ADSP Umbrella). https://dss.niagads.org/datasets/ng00067/ NIAGADS Data Sharing Service is needed to access the data.

The Cancer Genome Atlas data are available via dbGaP Study Accession (phs000178.v1.p1) and accessible via the National Cancer Institute Genomic Data Commons (https://gdc.cancer.gov/access-data/obtaining-access-controlled-data)An application to the

## ACKNOWLEDGMENTS

This work was supported in part by R01GM126249-03 (Bush, Crawford) and U01AG058654 (Haines, Bush, Farrer, Martin, Pericak-Vance).

The Alzheimer’s Disease Sequencing Project (ADSP) is comprised of two Alzheimer’s Disease (AD) genetics consortia and three National Human Genome Research Institute (NHGRI) funded Large Scale Sequencing and Analysis Centers (LSAC). The two AD genetics consortia are the Alzheimer’s Disease Genetics Consortium (ADGC), funded by NIA (U01 AG032984), and the Cohorts for Heart and Aging Research in Genomic Epidemiology (CHARGE) funded by NIA (R01 AG033193), the National Heart, Lung, and Blood Institute (NHLBI), other National Institute of Health (NIH) institutes and other foreign governmental and nongovernmental organizations. The Discovery Phase analysis of sequence data is supported through UF1AG047133 (to Drs. Farrer, Haines, Mayeux, Pericak-Vance, and Schellenberg); U01AG049505 to Dr. Seshadri; U01AG049506 to Dr. Boerwinkle; U01AG049507 to Dr. Wijsman; and U01AG049508 to Dr. Goate and the Discovery Extension Phase analysis is supported through U01AG052411 to Dr. Goate, U01AG052410 to Dr. Pericak-Vance and U01 AG052409 to Drs. Seshadri and Fornage. Data generation and harmonization in the Follow-up Phases is supported by U54AG052427 to Drs. Schellenberg and Wang. The ADGC cohorts include: Adult Changes in Thought (ACT supported by NIA grant U01AG006781 to Drs. Larson and Crane), the Alzheimer’s Disease Centers (ADC), the Chicago Health and Aging Project (CHAP), the Memory and Aging Project (MAP), Mayo Clinic (MAYO), Mayo Parkinson’s Disease controls, University of Miami, the Multi-Institutional Research in Alzheimer’s Genetic Epidemiology Study (MIRAGE), the National Cell Repository for Alzheimer’s Disease (NCRAD), the National Institute on Aging Late Onset Alzheimer’s Disease Family Study (NIA-LOAD), the Religious Orders Study (ROS), the Texas Alzheimer’s Research and Care Consortium (TARC), Vanderbilt University/Case Western Reserve University (VAN/CWRU), the Washington Heights-Inwood Columbia Aging Project (WHICAP supported by NIA grant RF1AG054023 to Dr. Mayeux) and the Washington University Sequencing Project (WUSP), the Columbia University Hispanic-Estudio Familiar de Influencia Genetica de Alzheimer (EFIGA supported by NIA grant RF1AG015473 to Dr. Mayeux), the University of Toronto (UT), and Genetic Differences (GD). Analysis of ADGC cohorts us supported by NIA grants R01AG048927 and RF1AG057519 to Dr. Farrer. Efforts of ADGC investigators were also supported by grants from the NIA (R03AG054936) and National Library of Medicine (R01LM012535). The CHARGE cohorts are supported in part by National Heart, Lung, and Blood Institute (NHLBI) infrastructure grant R01HL105756 (Psaty), RC2HL102419 (Boerwinkle) and the neurology working group is supported by the National Institute on Aging (NIA) R01 grant AG033193. The CHARGE cohorts participating in the ADSP include the following: Austrian Stroke Prevention Study (ASPS), ASPS-Family study, and the Prospective Dementia Registry-Austria (ASPS/PRODEM-Aus), the Atherosclerosis Risk in Communities (ARIC) Study, the Cardiovascular Health Study (CHS), the Erasmus Rucphen Family Study (ERF), the Framingham Heart Study (FHS), and the Rotterdam Study (RS). ASPS is funded by the Austrian Science Fond (FWF) grant number P20545-P05 and P13180 and the Medical University of Graz. The ASPS-Fam is funded by the Austrian Science Fund (FWF) project I904),the EU Joint Programme - Neurodegenerative Disease Research (JPND) in frame of the BRIDGET project (Austria, Ministry of Science) and the Medical University of Graz and the Steiermärkische Krankenanstalten Gesellschaft. PRODEM-Austria is supported by the Austrian Research Promotion agency (FFG) (Project No. 827462) and by the Austrian National Bank (Anniversary Fund, project 15435. ARIC research is carried out as a collaborativestudysupportedbyNHLBIcontracts (HHSN268201100005C, HHSN268201100006C, HHSN268201100007C, HHSN268201100008C, HHSN268201100009C, HHSN268201100010C, HHSN268201100011C, and HHSN268201100012C). Neurocognitive data in ARIC is collected by U01 2U01HL096812, 2U01HL096814, 2U01HL096899, 2U01HL096902, 2U01HL096917 from the NIH (NHLBI, NINDS, NIA and NIDCD), and with previous brain MRI examinations funded by R01-HL70825 from the NHLBI. CHS research was supported by contracts HHSN268201200036C, HHSN268200800007C, N01HC55222, N01HC85079, N01HC85080, N01HC85081, N01HC85082, N01HC85083, N01HC85086, and grants U01HL080295 and U01HL130114 from the NHLBI with additional contribution from the National Institute of Neurological Disorders and Stroke (NINDS). Additional support was provided by R01AG023629, R01AG15928, and R01AG20098 from the NIA. FHS research is supported by NHLBI contracts N01-HC-25195 and HHSN268201500001I. This study was also supported by additional grants from the NIA (R01s AG054076, AG049607 and AG033040 and NINDS (R01NS017950). The ERF study as a part of EUROSPAN (European Special Populations Research Network) was supported by European Commission FP6 STRP grant number 018947 (LSHG-CT-2006-01947) and also received funding from the European Community’s Seventh Framework Programme (FP7/2007-2013)/grant agreement HEALTH-F4-2007-201413 by the European Commission under the programme “Quality of Life and Management of the Living Resources” of 5th Framework Programme (no. QLG2-CT-2002-01254). High-throughput analysis of the ERF data was supported by a joint grant from the Netherlands Organization for Scientific Research and the Russian Foundation for Basic Research (NWO-RFBR 047.017.043). The Rotterdam Study is funded by Erasmus Medical Center and Erasmus University, Rotterdam, the Netherlands Organization for Health Research and Development (ZonMw), the Research Institute for Diseases in the Elderly (RIDE), the Ministry of Education, Culture and Science, the Ministry for Health, Welfare and Sports, the European Commission (DG XII), and the municipality of Rotterdam. Genetic data sets are also supported by the Netherlands Organization of Scientific Research NWO Investments (175.010.2005.011, 911-03-012), the Genetic Laboratory of the Department of Internal Medicine, Erasmus MC, the Research Institute for Diseases in the Elderly (014-93-015; RIDE2), and the Netherlands Genomics Initiative (NGI)/Netherlands Organization for Scientific Research (NWO) Netherlands Consortium for Healthy Aging (NCHA), project 050-060-810. All studies are grateful to their participants, faculty and staff. The content of these manuscripts is solely the responsibility of the authors and does not necessarily represent the official views of the National Institutes of Health or the U.S. Department of Health and Human Services. The ADES-FR study was funded by grants from the Clinical Research Hospital Program from the French Ministry of Health (GMAJ, PHRC, 2008/067), the CNR-MAJ, the JPND PERADES, the GENMED labex (LABEX GENMED ANR-10-LABX-0013), and the FP7 AgedBrainSysBio. Whole exome sequencing in the 3C-Dijon study was funded by the Fondation Leducq. This work was supported by the France Génomique National infrastructure, funded as part of the Investissements d’Avenir program managed by the Agence Nationale pour la Recherche (ANR-10-INBS-09), the Centre National de Recherche en Génomique Humaine, the National Foundation forAlzheimer’s disease and related disorders, the Institut Pasteur de Lille, Inserm, the Lille Métropole Communauté Urbaine council, and the French government’s LABEX (laboratory of excellence program investment for the future) DISTALZ grant (Development of Innovative Strategies for a Transdisciplinary approach to Alzheimer’s disease). The 3C Study supports are listed on the Study Website (www.three-city-study.com). The FinnAD Study at the University of Tampere was supported by The Academy of Finland: grants 286284 (T.L), Competitive State Research Financing of the Expert Responsibility area of Tampere University Hospitals (grant X51001); Juho Vainio Foundation; Paavo Nurmi Foundation; Finnish Foundation for Cardiovascular Research; Finnish Cultural Foundation; Tampere Tuberculosis Foundation; Yrjö Jahnsson Foundation; Signe and Ane Gyllenberg Foundation; and Diabetes Research Foundation of Finnish Diabetes Association. The FinnAD Study at the University of Eastern Finland was supported by the Academy of Finland grant 307866, the Sigrid Jusélius Foundation, and the Strategic Neuroscience Funding of the University of Eastern Finland. The three LSACs are: the Human Genome Sequencing Center at the Baylor College of Medicine (U54 HG003273), the Broad Institute Genome Center (U54HG003067), and the Washington University Genome Institute (U54HG003079).

Biological samples and associated phenotypic data used in primary data analyses were stored at Study Investigator institutions, and at the National Cell Repository for Alzheimer’s Disease (NCRAD, U24AG021886) at Indiana University funded by NIA. Associated Phenotypic Data used in primary and secondary data analyses were provided by Study Investigators, the NIA funded Alzheimer’s Disease Centers (ADCs), and the National Alzheimer’s Coordinating Center (NACC, U01AG016976) and the National Institute on Aging Genetics of Alzheimer’s Disease Data Storage Site (NIAGADS, U24AG041689) at the University of Pennsylvania, funded by NIA, and at the Database for Genotypes and Phenotypes (dbGaP) funded by NIH. This research was supported in part by the Intramural Research Program of the National Institutes of health, National Library of Medicine. Contributors to the Genetic Analysis Data included Study Investigators on projects that were individually funded by NIA, and other NIH institutes, and by private U.S. organizations, or foreign governmental or nongovernmental organizations. We also acknowledge the investigators who assembled and characterized participants of cohorts included in this study: Adult Changes in Thought: James D. Bowen, Paul K. Crane, Gail P. Jarvik, C. Dirk Keene, Eric B. Larson, W. William Lee, Wayne C. McCormick, Susan M. McCurry, Shubhabrata Mukherjee, Katie Rose Richmire Atherosclerosis Risk in Communities Study: Rebecca Gottesman, David Knopman, Thomas H. Mosley, B. Gwen Windham,

Austrian Stroke Prevention Study: Thomas Benke, Peter Dal-Bianco, Edith Hofer, Gerhard Ransmayr, Yasaman Saba

Cardiovascular Health Study: James T. Becker, Joshua C. Bis, Annette L. Fitzpatrick, M. Ilyas Kamboh, Lewis H. Kuller, WT Longstreth, Jr, Oscar L. Lopez, Bruce M. Psaty, Jerome I. Rotter, Chicago Health and Aging Project: Philip L. De Jager, Denis A. Evans

Erasmus Rucphen Family Study: Hieab H. Adams, Hata Comic, Albert Hofman, Peter J. Koudstaal, Fernando Rivadeneira, Andre G. Uitterlinden, Dina Voijnovic

Estudio Familiar de la Influencia Genetica en Alzheimer: Sandra Barral, Rafael Lantigua, Richard Mayeux, Martin Medrano, Dolly Reyes-Dumeyer, Badri Vardarajan

Framingham Heart Study: Alexa S. Beiser, Vincent Chouraki, Jayanadra J. Himali, Charles C. White

Genetic Differences: Duane Beekly, James Bowen, Walter A. Kukull, Eric B. Larson, Wayne McCormick, Gerard D. Schellenberg, Linda Teri

Mayo Clinic: Minerva M. Carrasquillo, Dennis W. Dickson, Nilufer Ertekin-Taner, Neill R. Graff-Radford, Joseph E. Parisi, Ronald C. Petersen, Steven G. Younkin

Mayo PD: Gary W. Beecham, Dennis W. Dickson, Ranjan Duara, Nilufer Ertekin-Taner, Tatiana M. Foroud, Neill R. Graff-Radford, Richard B. Lipton, Joseph E. Parisi, Ronald C. Petersen, Bill Scott, Jeffery M. Vance

Memory and Aging Project: David A. Bennett, Philip L. De Jager

Multi-Institutional Research in Alzheimer’s Genetic Epidemiology Study: Sanford Auerbach, Helan Chui, Jaeyoon Chung, L. Adrienne Cupples, Charles DeCarli, Ranjan Duara, Martin Farlow, Lindsay A. Farrer, Robert Friedland, Rodney C.P. Go, Robert C. Green, Patrick Griffith, John Growdon, Gyungah R. Jun, Walter Kukull, Alexander Kurz, Mark Logue, Kathryn L. Lunetta, Thomas Obisesan, Helen Petrovitch, Marwan Sabbagh, A. Dessa Sadovnick, Magda Tsolaki

National Cell Repository for Alzheimer’s Disease: Kelley M. Faber, Tatiana M. Foroud

National Institute on Aging (NIA) Late Onset Alzheimer’s Disease Family Study: David A. Bennett, Sarah Bertelsen, Thomas D. Bird, Bradley F. Boeve, Carlos Cruchaga, Kelley Faber, Martin Farlow, Tatiana M Foroud, Alison M Goate, Neill R. Graff-Radford, Richard Mayeux, Ruth Ottman, Dolly Reyes-Dumeyer, Roger Rosenberg, Daniel Schaid, Robert A Sweet, Giuseppe Tosto, Debby Tsuang, Badri Vardarajan

NIA Alzheimer Disease Centers: Erin Abner, Marilyn S. Albert, Roger L. Albin, Liana G. Apostolova, Sanjay Asthana, Craig S. Atwood, Lisa L. Barnes, Thomas G. Beach, David A. Bennett, Eileen H. Bigio, Thomas D. Bird, Deborah Blacker, Adam Boxer, James B. Brewer, James R. Burke, Jeffrey M. Burns, Joseph D. Buxbaum, Nigel J. Cairns, Chuanhai Cao, Cynthia M. Carlsson, Richard J. Caselli, Helena C. Chui, Carlos Cruchaga, Mony de Leon, Charles DeCarli, Malcolm Dick, Dennis W. Dickson, Nilufer Ertekin-Taner, David W. Fardo, Martin R. Farlow, Lindsay A. Farrer, Steven Ferris, Tatiana M. Foroud, Matthew P. Frosch, Douglas R. Galasko, Marla Gearing, David S. Geldmacher, Daniel H. Geschwind, Bernardino Ghetti, Carey Gleason, Alison M. Goate, Teresa Gomez-Isla, Thomas Grabowski, Neill R. Graff-Radford, John H. Growdon, Lawrence S. Honig, Ryan M. Huebinger, Matthew J. Huentelman, Christine M. Hulette, Bradley T. Hyman, Suman Jayadev, Lee-Way Jin, Sterling Johnson, M. Ilyas Kamboh, Anna Karydas, Jeffrey A. Kaye, C. Dirk Keene, Ronald Kim, Neil W Kowall, Joel H. Kramer, Frank M. LaFerla, James J. Lah, Allan I. Levey, Ge Li, Andrew P. Lieberman, Oscar L. Lopez, Constantine G. Lyketsos, Daniel C. Marson, Ann C. McKee, Marsel Mesulam, Jesse Mez, Bruce L. Miller, Carol A. Miller, Abhay Moghekar, John C. Morris, John M. Olichney, Joseph E. Parisi, Henry L. Paulson, Elaine Peskind, Ronald C. Petersen, Aimee Pierce, Wayne W. Poon, Luigi Puglielli, Joseph F. Quinn, Ashok Raj, Murray Raskind, Eric M. Reiman, Barry Reisberg, Robert A. Rissman, Erik D. Roberson, Howard J. Rosen, Roger N. Rosenberg, Martin Sadowski, Mark A. Sager, David P. Salmon, Mary Sano, Andrew J. Saykin, Julie A. Schneider, Lon S. Schneider, William W. Seeley, Scott Small, Amanda G. Smith, Robert A. Stern, Russell H. Swerdlow, Rudolph E. Tanzi, Sarah E Tomaszewski Farias, John Q. Trojanowski, Juan C. Troncoso, Debby W. Tsuang, Vivianna M. Van Deerlin, Linda J. Van Eldik, Harry V. Vinters, Jean Paul Vonsattel, Jen Chyong Wang, Sandra Weintraub, Kathleen A. Welsh-Bohmer, Shawn Westaway, Thomas S. Wingo, Thomas Wisniewski, David A. Wolk, Randall L. Woltjer, Steven G. Younkin, Lei Yu, Chang-En Yu

Religious Orders Study: David A. Bennett, Philip L. De Jager

Rotterdam Study: Kamran Ikram, Frank J Wolters

Texas Alzheimer’s Research and Care Consortium: Perrie Adams, Alyssa Aguirre, Lisa Alvarez, Gayle Ayres, Robert C. Barber, John Bertelson, Sarah Brisebois, Scott Chasse, Munro Culum, Eveleen Darby, John C. DeToledo, Thomas J. Fairchild, James R. Hall, John Hart, Michelle Hernandez, Ryan Huebinger, Leigh Johnson, Kim Johnson, Aisha Khaleeq, Janice Knebl, Laura J. Lacritz, Douglas Mains, Paul Massman, Trung Nguyen, Sid O’Bryant, Marcia Ory, Raymond Palmer, Valory Pavlik, David Paydarfar, Victoria Perez, Marsha Polk, Mary Quiceno, Joan S. Reisch, Monica Rodriguear, Roger Rosenberg, Donald R. Royall, Janet Smith, Alan Stevens, Jeffrey L. Tilson, April Wiechmann, Kirk C. Wilhelmsen, Benjamin Williams, Henrick Wilms, Martin Woon

University of Miami: Larry D Adams, Gary W. Beecham, Regina M Carney, Katrina Celis, Michael L Cuccaro, Kara L. Hamilton-Nelson, James Jaworski, Brian W. Kunkle, Eden R. Martin, Margaret A. Pericak-Vance, Farid Rajabli, Michael Schmidt, Jeffery M Vance

University of Toronto: Ekaterina Rogaeva, Peter St. George-Hyslop

University of Washington Families: Thomas D. Bird, Olena Korvatska, Wendy Raskind, Chang-En Yu

Vanderbilt University: John H. Dougherty, Harry E. Gwirtsman, Jonathan L. Haines

Washington Heights-Inwood Columbia Aging Project: Adam Brickman, Rafael Lantigua, Jennifer Manly, Richard Mayeux, Christiane Reitz, Nicole Schupf, Yaakov Stern, Giuseppe Tosto, Badri Vardarajan

## REFERENCE

Akiyama, H., Nishimura, T., Kondo, H., Ikeda, K., Hayashi, Y., and McGeer, P. L. (1994). Expression of the receptor for macrophage colony stimulating factor by brain microglia and its upregulation in brains of patients with Alzheimer’s disease and amyotrophic lateral sclerosis. Brain research, 639(1), 171–174.

Beecham, G.W., Bis, J.C., Martin, E.R., Choi, S.-H., DeStefano, A.L., van Duijn, C.M., Fornage, M., Gabriel, S.B., Koboldt, D.C., Larson, D.E., et al. (2017). The Alzheimer’s Disease Sequencing Project: Study design and sample selection. Neurol Genet 3, e194.

Bis, J.C., Jian, X., Kunkle, B.W., Chen, Y., Hamilton-Nelson, K.L., Bush, W.S., Salerno, W.J., Lancour, D., Ma, Y., Renton, A.E., et al. (2020). Whole exome sequencing study identifies novel rare and common Alzheimer’s-Associated variants involved in immune response and transcriptional regulation. Mol Psychiatry 25, 1859–1875.

Butkiewicz, M., Blue, E.E., Leung, Y.Y., Jian, X., Marcora, E., Renton, A.E., Kuzma, A., Wang, L.-S., Koboldt, D.C., Haines, J.L., et al. (2018). Functional annotation of genomic variants in studies of late-onset Alzheimer’s disease. Bioinformatics 34, 2724–2731.

Dana, J.M., Gutmanas, A., Tyagi, N., Qi, G., O’Donovan, C., Martin, M., and Velankar, S. (2019). SIFTS: updated Structure Integration with Function, Taxonomy and Sequences resource allows 40-fold increase in coverage of structure-based annotations for proteins. Nucleic Acids Research 47, D482–D489.

Guerreiro, R., Wojtas, A., Bras, J., Carrasquillo, M., Rogaeva, E., Majounie, E., Cruchaga, C., Sassi, C., Kauwe, J. S. K., et al. (2013). TREM2 variants in Alzheimer’s disease. New England Journal of Medicine, 368, 117–127.

Henikoff, S., and Henikoff, J.G. (1992). Amino acid substitution matrices from protein blocks. Proc Natl Acad Sci U S A 89, 10915–10919.

Jeong, D.G., Cho, Y.H., Yoon, T.S., Kim, J.H., Son, J.H., Ryu, S.E., and Kim, S.J. (2006). Structure of human DSP18, a member of the dual-specificity protein tyrosine phosphatase family. Acta Crystallogr D Biol Crystallogr 62, 582–588.

Kamburov, A., Lawrence, M.S., Polak, P., Leshchiner, I., Lage, K., Golub, T.R., Lander, E.S., and Getz, G. (2015). Comprehensive assessment of cancer missense mutation clustering in protein structures. Proc Natl Acad Sci USA 112, E5486–E5495.

Karczewski, K.J., Francioli, L.C., Tiao, G., Cummings, B.B., Alföldi, J., Wang, Q., Collins, R.L., Laricchia, K.M., Ganna, A., Birnbaum, D.P., et al. (2020). The mutational constraint spectrum quantified from variation in 141,456 humans. Nature 581, 434–443.

Korvatska, O., Leverenz, J.B., Jayadev, S., McMillan, P., Kurtz, I., Guo, X., Rumbaugh, M., Matsushita, M., Girirajan, S., Dorschner, M.O., et al. (2015). R47H Variant of TREM2 Associated With Alzheimer Disease in a Large Late-Onset Family: Clinical, Genetic, and Neuropathological Study. JAMA Neurol 72, 920–927.

Li, B., and Leal, S. M. (2008). Methods for detecting associations with rare variants for common diseases: application to analysis of sequence data. The American Journal of Human Genetics, 83, 311–321.

Leung, Y.Y., Valladares, O., Chou, Y.-F., Lin, H.-J., Kuzma, A.B., Cantwell, L., Qu, L., Gangadharan, P., Salerno, W.J., Schellenberg, G.D., et al. (2019). VCPA: genomic variant calling pipeline and data management tool for Alzheimer’s Disease Sequencing Project. Bioinformatics 35, 1768–1770.

Mashl, R.J., Wu, Y., Ritter, D.I., Wang, J., Oh, C., Paczkowska, M., Reynolds, S., Wyczalkowski, M.A., Oak, N., Scott, A.D., et al. (2018). Pathogenic Germline Variants in 10,389 Adult Cancers. Cell 173, 355–370.e14.

Marceau West, R., Lu, W., Rotroff, D.M., Kuenemann, M.A., Chang, S.-M., Wu, M.C., Wagner, M.J., Buse, J.B., Motsinger-Reif, A.A., Fourches, D., et al. (2019). Identifying individual risk rare variants using protein structure guided local tests (POINT). PLoS Comput Biol 15, e1006722.

Monticelli, L., Kandasamy, S.K., Periole, X., Larson, R.G., Tieleman, D.P., and Marrink, S.-J. (2008). The MARTINI Coarse-Grained Force Field: Extension to Proteins. J. Chem. Theory Comput. 4, 819–834.

Ryu, J., and Lee, D.H. (2018). Dual-specificity phosphatase 18 modulates the SUMOylation and aggregation of Ataxin-1. Biochem Biophys Res Commun 502, 389–396.

Senior, A.W., Evans, R., Jumper, J., Kirkpatrick, J., Sifre, L., Green, T., Qin, C., Žídek, A., Nelson, A.W.R., Bridgland, A., et al. (2020). Improved protein structure prediction using potentials from deep learning. Nature 577, 706–710.

Sivley, R.M., Sheehan, J.H., Kropski, J.A., Cogan, J., Blackwell, T.S., Phillips, J.A., Bush, W.S., Meiler, J., and Capra, J.A. (2018). Three-dimensional spatial analysis of missense variants in RTEL1 identifies pathogenic variants in patients with Familial Interstitial Pneumonia. BMC Bioinformatics 19, 18.

Somody, J.C., MacKinnon, S.S., and Windemuth, A. (2017). Structural coverage of the proteome for pharmaceutical applications. Drug Discov Today 22, 1792–1799.

Stanley, E. R., and Chitu, V. (2014). CSF-1 receptor signaling in myeloid cells. Cold Spring Harbor perspectives in biology, 6, a021857.

Suh, J., Romano, D.M., Nitschke, L., Herrick, S.P., DiMarzio, B.A., Dzhala, V., Bae, J.-S., Oram, M.K., Zheng, Y., Hooli, B., et al. (2019). Loss of Ataxin-1 Potentiates Alzheimer’s Pathogenesis by Elevating Cerebral BACE1 Transcription. Cell 178, 1159–1175.e17.

Taliun, D., Harris, D. N., Kessler, M. D., Carlson, J., Szpiech, Z. A., Torres, R., Taliun, S. A., Corvelo, A., Gogarten, S. M., et al. (2021). Sequencing of 53,831 diverse genomes from the NHLBI TOPMed Program. Nature, 590, 290–299.

Tang, Z.-Z., Sliwoski, G.R., Chen, G., Jin, B., Bush, W.S., Li, B., and Capra, J.A. (2020). PSCAN: Spatial scan tests guided by protein structures improve complex disease gene discovery and signal variant detection. Genome Biol 21, 217.

Tokheim, C., Bhattacharya, R., Niknafs, N., Gygax, D.M., Kim, R., Ryan, M., Masica, D.L., and Karchin, R. (2016). Exome-Scale Discovery of Hotspot Mutation Regions in Human Cancer Using 3D Protein Structure. Cancer Res 76, 3719–3731.

Vardarajan, B. N., Zhang, Y., Lee, J. H., Cheng, R., Bohm, C., Ghani, M., Reitz, C., Reyes-Dumeyer, D., Shen, Y., et al. (2015). Coding mutations in SORL 1 and A lzheimer disease. Annals of neurology, 77, 215–227.

Wu, M.C., Lee, S., Cai, T., Li, Y., Boehnke, M., and Lin, X. (2011). Rare-Variant Association Testing for Sequencing Data with the Sequence Kernel Association Test. The American Journal of Human Genetics 89, 82–93.

Wang, K. (2016). Boosting the power of the sequence kernel association test by properly estimating its null distribution. The American Journal of Human Genetics, 99, 104–114.

